# Control of Leaf Width by the APC/C^TAD1^-WL1-NAL1 Pathway in Rice

**DOI:** 10.1101/2022.01.10.475683

**Authors:** Jing You, Wenwen Xiao, Yue Zhou, Li Ye, Peng Yu, Guoling Yu, Xinfang Zhang, Zhifeng He, Yan Xiang, Xianchun Sang, Yunfeng Li, Fangming Zhao, Yinghua Ling, Guanghua He, Ting Zhang

## Abstract

Leaf morphology is one of the most important features of the ideal plant architecture. However, the genetic and molecular mechanisms controlling leaf morphology in crops remain largely unknown, despite their central importance. Here we demonstrate that the APC/C^TAD1^-WL1-NAL1 pathway regulates leaf width in rice, and mutation of *WL1* leads to width leaf variation. WL1 interacts with TAD1 and is degraded by APC/C^TAD1^, with the loss of *TAD1* function resulting in narrow leaves. The WL1 protein directly binds to the regulatory region of *NAL1* and recruits the corepressor TOPLESS-RELATED PROTEIN to inhibit *NAL1* expression by down-regulating the level of histone acetylation of chromatin. Furthermore, biochemical and genetic analyses revealed that TAD1, WL1, and NAL1 function in a common pathway to control leaf width. Our study establishes an important framework for the APC/C^TAD1^-WL1-NAL1 pathway-mediated control of leaf width in rice and introduces novel perspectives for using this regulatory pathway for improving crop plant architecture.

## Introduction

Leaves are important for photosynthesis and leaf width is one of the most critical components of the ideal plant architecture. Appropriate leaf width is of great biological significance for improving light energy absorption and conversion efficiency(Ort et al., 2015). Optimal leaf width is a key determinant of rice yield as well as an important research goal for developing ideal super-high-yielding crop plants. Therefore, in-depth studies on the mechanisms of regulation of leaf width are important theoretically and practically for improving rice yield.

Leaves develop from the shoot apical meristems. Specialized rapidly dividing cells on the periphery of the shoot apical meristem develop into leaf primordia (Fleming, 2005; Moon and Hake, 2011). Afterwards, the cells alter their plane of division and establish three-dimensional axes along the proximal/distal, adaxial/abaxial, and medial/lateral axes (Moon and Hake, 2011). Auxin is involved in the regulation of leaf width, with two narrow leaf genes, *NAL1* and *NAL7,* influence auxin polar transport and synthesis, respectively. *NAL1* encodes a plant-specific protein with largely unknown biochemical functions. Mutation of *NAL1* leads to a significant reduction in polar auxin transport capacity and thereby alters the distribution pattern of vascular tissues and reduces leaf width (Qi et al., 2008a; Chen et al., 2012). *NAL7* encodes a flavin-containing monooxygenase which exhibits sequence homology with YUCCA and is involved in auxin biosynthesis. The *nal7* mutant has a reduced auxin content and shows a narrow leaf phenotype (Fujino et al., 2008a). Leaf width in rice is also closely related to leaf vascular development. *NAL2* and *NAL3* are paralogs that encode identical OsWOX3A transcriptional activators. The numbers of small veins between adjacent large veins in the *nal2/nal3* mutant are remarkably reduced, and the arrangement of xylem and phloem is abnormal, resulting in narrow leaves. The phenotypes of these mutants indicate that *NAL2/NAL3* has a significant regulatory effect on the development of vascular bundles (Cho et al., 2013; Ishiwata et al., 2013). *NRL1* encodes the cellulose synthase-like protein D4 (OsCslD4). Mutation of *NRL1* results in a decreased number of longitudinal veins in leaves, producing phenotypes with reduced leaf width(Li et al., 2009; Yoshikawa et al., 2013). Additionally, leaf width is also related to polarity. *SLL1* encodes a SHAQKYF class MYB transcription factor belonging to the KAN family. Mutation of *SLL1* results in narrowed and abaxialized leaves (Zhang et al., 2009). *SRL2* encodes a newly identified plant-specific protein of unknown biochemical function and the *srl2* mutant has narrow incurved leaves due to the defective development of abaxial sclerenchymatous cells (Ma et al., 2017).

The Cys-2/His-2-type (C2H2) zinc finger protein is an important class of eukaryotic transcription factors, which has been implicated in different cellular mechanisms involved in developmental processes and in plant responses to stress (Sun et al., 2010). *DST* (*DROUGHTAND SALT TOLERANCE*) encodes a novel C2H2 zinc finger transcription factor that contains a highly conserved C2H2-type zinc finger domain and exhibits transcriptional activation function. *DST* plays an important role in the responses of plants to stress. *DST* negatively regulates stomatal closure by directly modulating genes related to H_2_O_2_ homeostasis through a novel signal transduction pathway for *DST*-mediated H_2_O_2_-induced stomatal closure. Loss of *DST* function promotes stomatal closure and reduces stomatal density, resulting in enhanced drought and salt tolerance in rice (Huang et al., 2009). *DST* also regulates drought and salt tolerance through interacting with the co-activator DST Co-activator 1 (DCA1) to regulate the expression of a peroxidase 24 precursor (*Prx 24*), a gene that encodes an H_2_O_2_ scavenger (Cui et al., 2015). DST can also regulate the expression of a leucine-rich repeat (LRR)-RLK gene named *Leaf Panicle 2* (*LP2*) under drought stress (Wu et al., 2015). Additionally, *DST* plays an important role in the biological development where it positively and directly regulates the expression of *OsCKX2* leading to increased CK accumulation in the shoot apical meristem (SAM) and enhanced seed production. Mutants of *DST* show increased meristematic activity, enhanced panicle branching and a consequent increase in grain number per panicle (Li et al., 2013). Further studies showed that DST interacts with OsMPK6 and is phosphorylated by OsMPK6, thus enhancing its capacity to activate the transcription of *OsCKX2* (Guo et al., 2020). Generally, *DST*plays a dual role in regulating stress and developmental responses. However, how *DST* participates in the regulation of leaf width is still largely unknown.

In this study, we isolated a *wide-leaf 1* (*wl1*) mutant, which exhibited an increased leaf width. We report that the *wl1* phenotype was caused by a novel allelic mutation in the *DST* gene, which encodes a zinc finger protein. Although a function of *WL1/DST* in drought and salt tolerance in rice has been reported (Huang et al., 2009; Li et al., 2013; Cui et al., 2015), the role of *WL1/DST* in controlling leaf width has not been studied in detail. We further demonstrate that the APC/C^TAD1^-WL1-NAL1 signaling pathway positively regulates leaf width in rice where mutation of *WL1* resulted in variation of leaf width. WL1 interacted with TAD1 and acted as a degradable substrate of APC/C^TAD1^. Loss of *TAD1* function resulted in narrow leaf. WL1 protein was able to bind to the regulatory region of *NAL1* directly and then recruit the corepressor TOPLESS-RELATED PROTEIN to repress the expression of *NAL1* by down-regulating the histone acetylation levels of chromatin. Loss of *NAL1* function also resulted in narrow leaves. Further biochemical and genetic analyses revealed that TAD1, WL1, and NAL1 functioned in a common pathway to control leaf width. Taken together, our study establishes an important genetic and molecular framework for the APC/C^TAD1^-WL1-NAL1 pathway-mediated control of leaf width in rice. This suggests that this pathway is crucial for regulation of leaf architecture in crop plants.

## Results

### Phenotypic identification of the *wl1* mutant

Compared with wild-type (WT) Xinong 1B, the most remarkable feature of the *wl1* mutant was an increased leaf width throughout the growth season (Figure 1A-F). The leaf width and length of the WT and *wl1* were measured at the same time points in the growth period. The leaf width of *wl1* was significantly higher than that of the WT throughout the growth period (Figure 1G). However, the leaf length of *wl1* did not vary significantly from that of the WT (Figure 1H).

**Figure 1.**
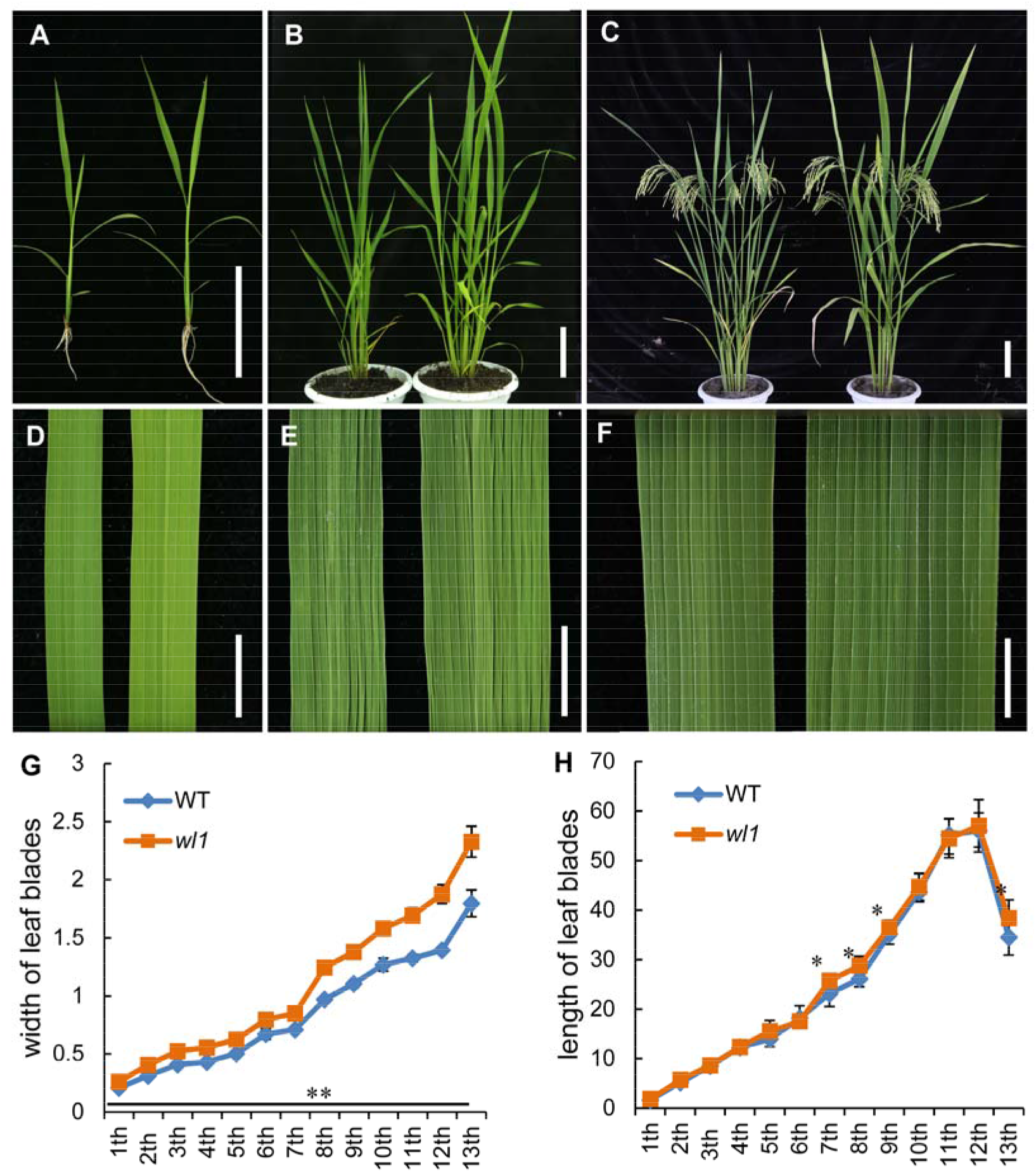
Phenotypic characteristics of the *wl1* mutant. (A–F) Phenotypes of the WT (left) and *wl1* (right) at the seedling stage (A, D), tiller stage (B, E), and heading stage (C, F). (G–H) Leaf blade width (G) and length (H) of the WT and *wl1* plants for the 1^st^–14^th^ leaves produced across the total growth period. *p* values were determined using t-test as compared with Xinong 1B. ** represents *p*≤0.01. Bar=10 cm in A-C and 1 cm in D-F.

To investigate the developmental sequence leading to the wide leaf phenotype in *wl1*, histological analyses of the shoot apical meristems (SAM), leaf primordia and leaf blades were performed. We found that the size and shape of the SAM of *wl1* were not significantly different from those of the WT (Figure 2 A, B, I). However, the area and width of P1, P2 and P3 leaf primordia in the *wl1* mutant were significantly greater than those in the WT (Figure 2 C, D, J, L). Subsequent histological analysis of the fifth mature leaves of the tillers revealed that the *wl1* leaves were significantly wider than those of the wild-type (Figure 2 E, F). Notably, the distance between the small vascular bundles in *wl1* increased by 22.1% compared to the WT (Figure 2K). The numbers of small and large vascular bundles were significantly increased compared to the WT throughout the whole growth season (Figure 2 G, H). There were no visible changes in the shape of the midrib or the large and small vascular bundles (Supplemental Figure 1). Therefore, the distinctly wide leaf of *wl1* originated at the developmental stage of leaf primordium and was mainly dependent on the number of vascular bundles and the distances between small vascular bundles.

**Figure 2.**
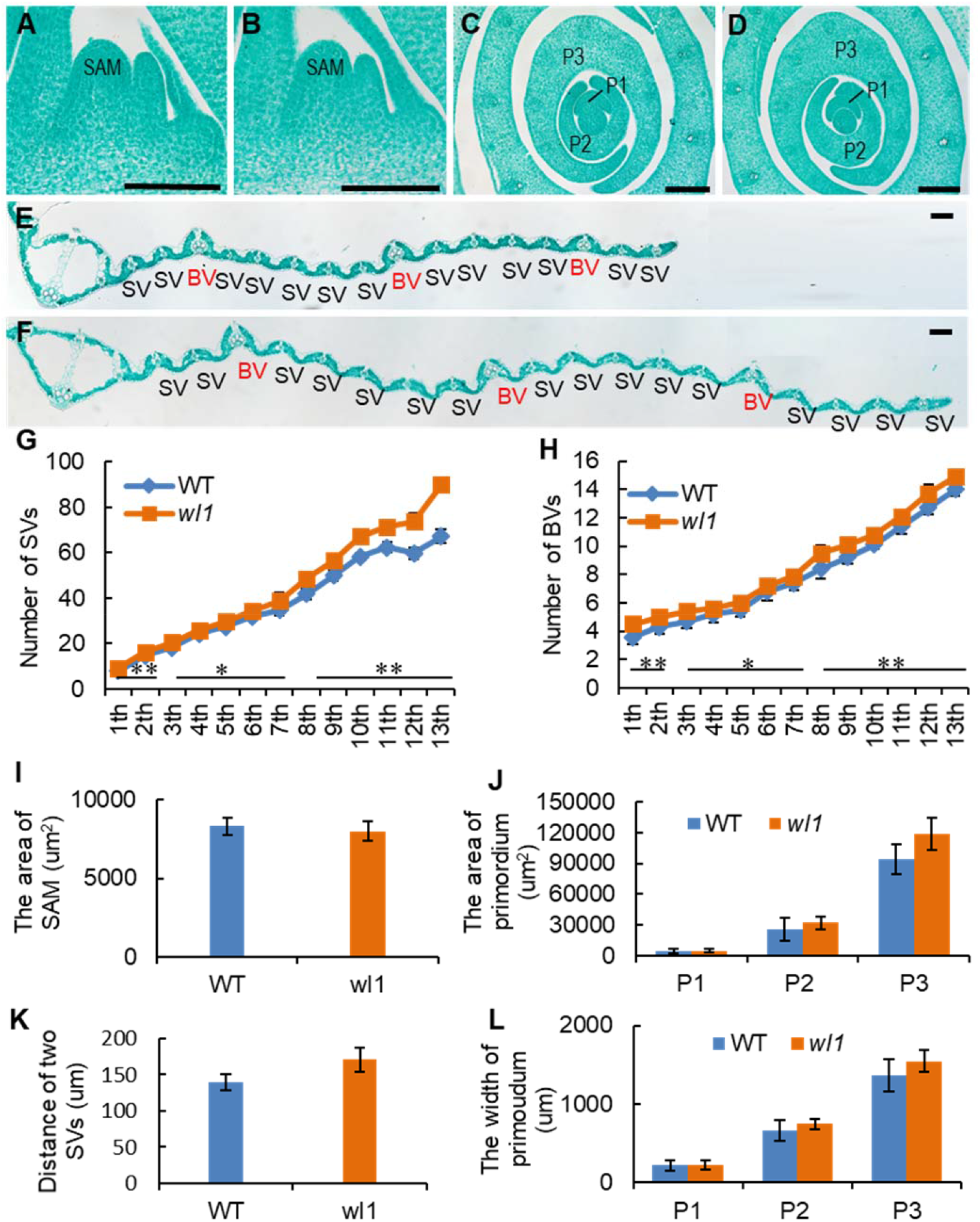
Histological analysis of the wild-type (WT) and *wl1* mutant. (A–B) Longitudinal sections of the shoot apex of WT (A) and *wl1* (B). (C–D) Transverse sections of the shoot apex of WT (**c**) and *wl1* (D). (E-F) Transverse sections of the leaf blade of WT (**e**) and *wl1* (F). (G–L) Comparisons of the number of small vascular bundles (SVs) (G) and large vascular bundles (BVs). (H), The area of SAM, (I), the area of primordium (J), the distance of two SVs (K), and (L) the width of primordium. SAM, Shoot apical meristem; P2, the second primordium; P3, the third primordium. Bar=100 μm.

### Molecular cloning and identification of the *WL1* gene

A map-based cloning strategy was adopted to isolate the *WL1* gene. Genetic analysis of the F_2_ progeny revealed that the segregation ratio of the WT and individual mutant plants showed a good fit to 3:1 (544 of 1,648 were mutant individuals; *χ*^2^ = 0.037 < *χ*^2^_0.05_ = 3.84), indicating that a single recessive gene controls the *wl1* trait. Using the 544 mutant plants in the F_2_ population, we narrowed the location of the *WL1* gene to a 53.31-kb region between the markers ID56 and ID65 on the long arm of chromosome 3. In this region, there were six annotated genes based on the Michigan State University (MSU) Rice Genome Annotation Project (Figure 3A). In the *wl1* mutant, a single-nucleotide substitution from C to T was identified in the annotated gene *LOC_Os03g57240*. Such mutation converted a glutamine codon into a termination codon resulting in premature termination of protein translation (Figure 3A). *WL1* is an allelic gene of *DST*, which encodes a C2H2 zinc finger protein and regulates responses to drought and salt stress, as well as the grain number per panicle in rice (Huang et al., 2009; Li et al., 2013). The WL1 protein was localized to the nucleus (Supplemental Figure 2) and the homologous genes of *WL1* were found to be abundant and highly conserved in monocotyledons and dicotyledons (Supplemental Figure 3).

**Figure 3.**
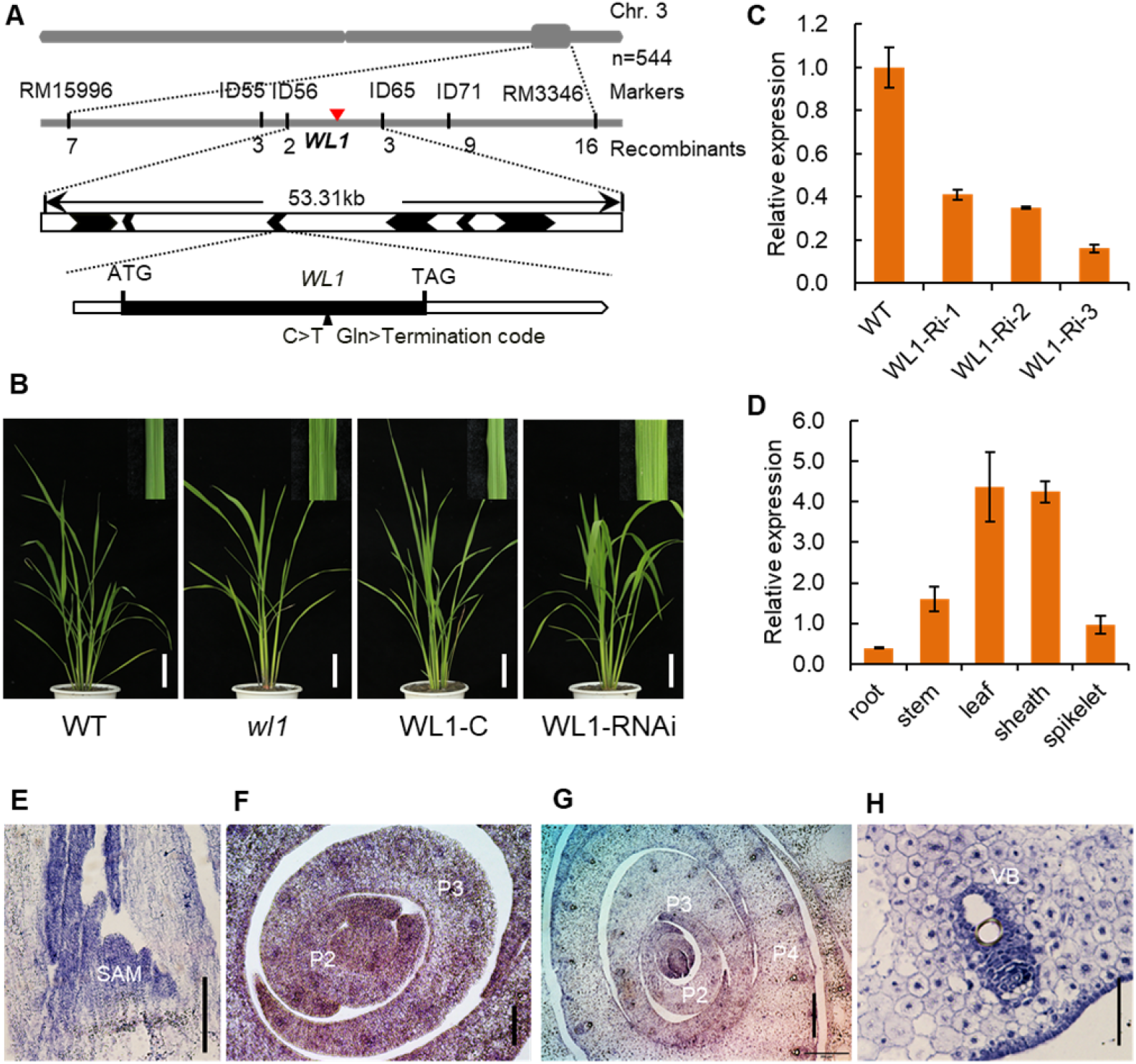
Map-Based cloning and expression patterns of *WL1*. (A) Mapping of *WL1* locus and the molecular lesions in the *Wl1* mutant. (B) Comparison of plants and leaf width of the WT, *wl1*, and the transgenic lines WL1-C (complementation), RNAi (RNA interference). (C) Expression analysis of *WL1* in leaves of the WT and the three RNAi lines by qRT-PCR. (D) Expression analysis of *WL1* in the roots, stems, leaves, leaf sheaths and spikelets of the WT. (E-H) Expression analysis of *WL1* in SAMs and leaf primordia of the WT by *in situ* hybridization. *p* values were determined using t-test as compared with Xinong 1B. SAM, shoot apical meristem; P1, the first primordium; P2, the second primordium; P3, the third primordium; VB, vascular bundle. Bars=10 cm in B; 200 μm in E-G; 50 μm in H.

To verify whether the mutation of *LOC_Os03g57240* is linked to the mutant phenotype, we performed a complementation experiment by introducing a 3776-bp WT DNA fragment containing *LOC_Os03g57240* into the *wl1* mutant. The mutant phenotype was rescued entirely in the transgenic plants (Figure 3B). Next, we performed RNAi suppression test to silence *WL1* in the WT plants. In the transgenic plants, the quantity of *WL1* transcripts was greatly reduced (Figure 3C) and the leaf width was rendered similar to that of the *wl1* mutant (Figure 3B). Overall, these results confirmed that *LOC_Os03g57240/DST* corresponds to the *WL1* gene.

The expression of *WL1* was investigated using quantitative reverse transcription-PCR (qRT-PCR) analysis. *WL1* was expressed in various tissues including the roots, stems, leaves, leaf sheaths and spikelets (Figure 3D). For a more detailed analysis of the expression pattern of *WL1*, *in situ* hybridization was conducted (Figure 3E-H). Signals of *WL1* were mainly concentrated in the SAM, leaf initiation site (P0) and leaf primordium (Figure 3E-H). Strong signals were observed in the SAM (Figure 3E). However, the signals in the P2 primordia were strong but scattered (Figure 3F). In the P3 primordia, *WL1* signals were concentrated at the margin of the primordium and the procambial cells (Figure 3E), potentially indicating the sites of differentiation of the procambial cells from fundamental cells. In the P4 primordia, *wL1* signals were largely restricted to the vascular bundles, with no preference for xylem or phloem (Figure 3 G, H). These results suggested that *wL1* may be involved in leaf development.

### WL1 Interacts with TAD1

To analyze the molecular mechanism of *WL1*-mediated regulation of leaf width, we first searched for proteins that may interact with the WL1 protein to regulate grain development. A yeast two-hybrid assay showed that WL1 interacted with TAD1 (Figure 4A), a co-activator of the anaphase-promoting complex (APC/C), which is a multi-subunit E3 ligase that regulates the growth of rice tillers and roots (Lin et al., 2012; Xu et al., 2012; Lin et al., 2020). The interaction between WL1 and TAD1 was verified by an in vitro pull-down assay. WL1 was fused to glutathione-S-transferase (GST-WL1) and His-tagged TAD1 (His-TAD1) and subsequently expressed in *Escherichia coli*. GST-WL1, but not GST, was able to pull down His-TAD1 (Figure 4B). Consistent results were obtained from the BiFC assay in the leaf epidermal cells of *Nicotiana benthamiana in vivo*. WL1 was fused to the C-terminal half of YFP (WL1-cYFP) and TAD1 was fused to the N-terminal half of YFP (TAD1-nYFP). The YFP signal was mainly restricted to the nuclei when WL1-cYFP was co-transformed with TAD1-nYFP (Figure 4C). These results confirmed the direct interaction between WL1 and TAD1 *in vitro* and *in vivo*. Furthermore, qRT-PCR results showed that *TAD1* was also expressed in various tissues including the roots, stems, leaves, leaf sheaths, and spikelets (Figure 4D). Thus, the expression pattern of *TAD1* was consistent with that of *WL1*, suggesting that WL1 interacts with TAD1 to regulate the leaf width.

**Figure 4.**
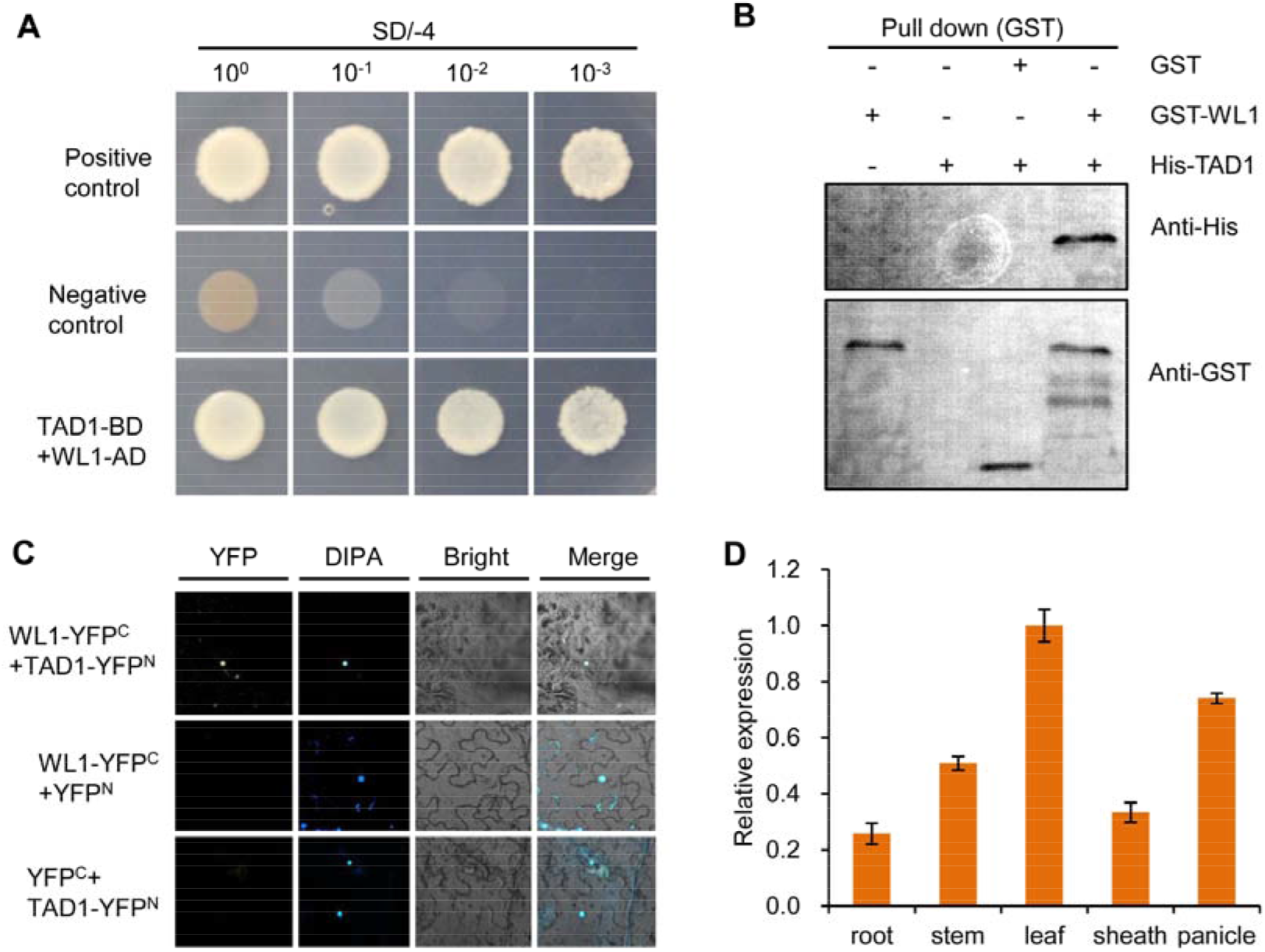
WL1 Interacts with TAD1. (A) WL1 interacts with TAD1 in yeast cells as shown by Y2H assay. (B) WL1 interacts with TAD1 *in vitro* as detected by pull-down assay. (C) WL1 interacts with TAD1 in the nuclei of *N. benthamiana* leaf cells as measured by the BiFC assay. (D) Expression analysis of *TAD1* in the roots, stems, leaves, leaf sheaths and spikelets of the WT.

### WL1 acts as a degradable substrate for APC/C^TAD1^

*TAD1* encodes a rice homologue of Cdh1 that functions as an activator of the anaphase-promoting complex/cyclosome (APC/C) (Lin et al., 2012). Based on this information, we carried out a series of biochemical experiments to examine whether the APC/C^TAD1^ complex mediates the degradation of WL1 via the ubiquitin-26S proteasome pathway. An *in vitro* ubiquitination assay demonstrated that WL1 was poly-ubiquitinated more efficiently by the total protein extracts of the WT than by those of the *tad1* plants (Figure 5A), indicating that TAD1 was required for ubiquitination of WL1. A cell-free degradation study showed that the purified recombinant GST-WL1 protein but not the GST control was rapidly degraded by the total protein extracts of the WT, but its degradation was compromised in the *tad1* mutant extracts (Figure 5B). Moreover, MG132, a specific inhibitor of the 26S proteasome, blocked the degradation of GST-WL1 in total protein extracts (Figure 5B).

**Figure 5.**
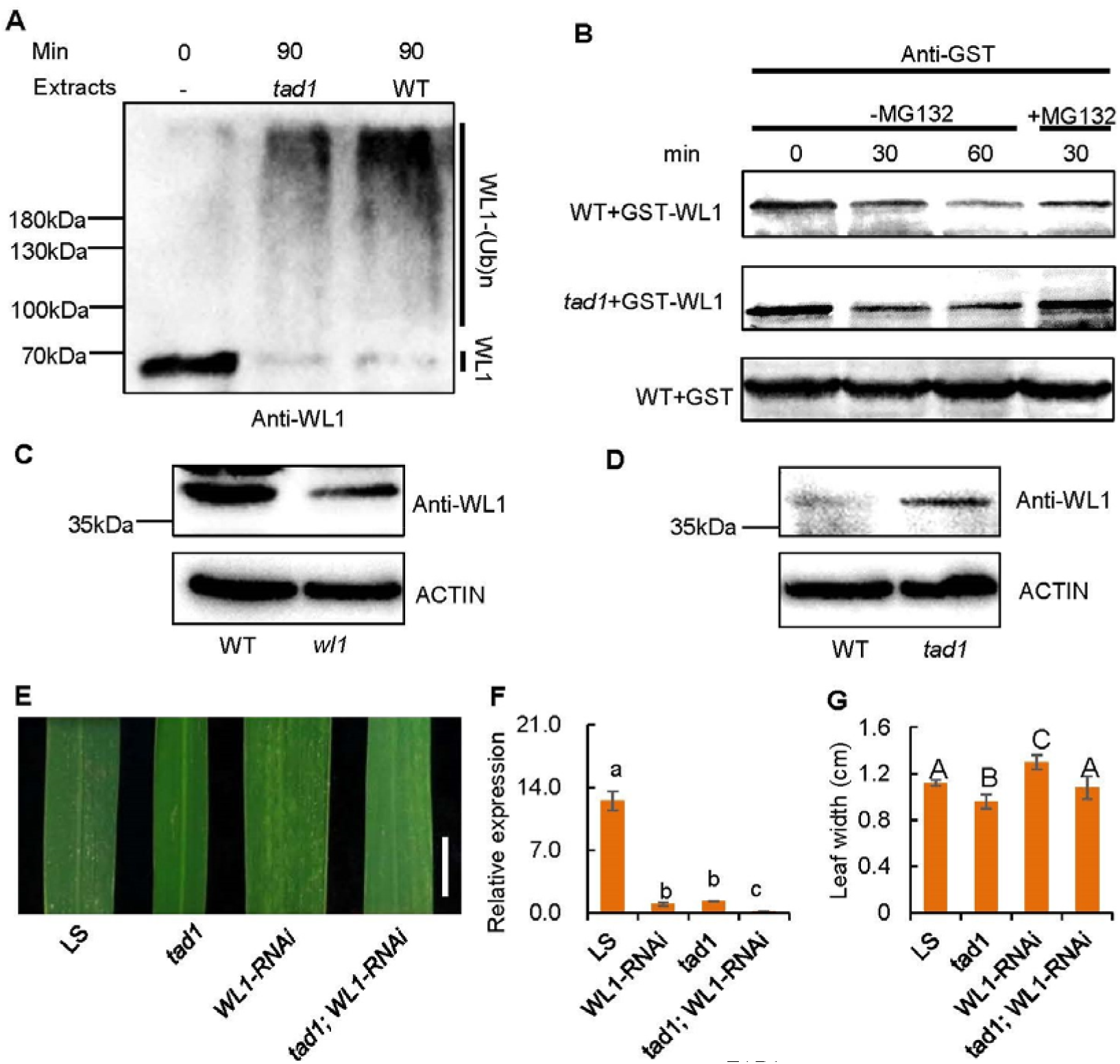
WL1 acts as a degradable substrate for APC/C^TAD1^. (A) *In vitro* ubiquitination assay showing that WL1 was ubiquitinated by protein extracts of the WT but not by those of *tad1* plants. (B) Cell-free degradation assay showing that GST-WL1, but not GST, was degraded in the protein extracts of the WT after 60 min of incubation, whereas GST-WL1 remained relatively stable in the extract of *tad1*. The degradation of GST-WL1 by protein extracts was inhibited in the presence of 40 μM MG132 (+MG132). (C) Western blot analysis showed that WL1 protein accumulated at lower concentrations in the *wl1* leaves compared to the WT (top). Actin strips showing approximately equal loading of total proteins (bottom). (D) Western blot analysis showed that WL1 protein accumulated to higher concentrations in the *tad1* leaves as compared to the WT (top). Actin strips showed approximately equal loading of total proteins (bottom). (E) Leaves of LS, tad1, WL1-RNAi, and *tad1*; WL1-RNAi. (F) Expression of *WL1* in LS, *tad1*, WL1-RNAi, and *tad1*; WL1-RNAi. (G) Leaf width of LS, *tad1*, WL1-RNAi and *tad1*; WL1-RNAi. *p* value was determined using t-test as compared with LS. Lowercase letters indicate significant differences (*p*<0.05). Scale bar=1 cm.

Additionally, WL1 protein and mRNA levels were remarkably lower in *wl1* plants but higher in *tad1* ones as compared to the WT (Figure 5C, D) (Supplemental Figure 4). Taken together, these results support the hypothesis that WL1 functions as an authentic degradable substrate for APC/C^TAD1^ to regulate leaf width in rice.

To further resolve the genetic interaction between WL1 and TAD1, we analyzed the leaf width of the *tad1* mutant. As hypothesized, the *tad1* mutant showed a narrow leaf phenotype where its leaf width was decreased significantly by 14.3% compared to that of the WT (LS) (Figure 5E, G). Furthermore, we generated RNAi lines of *WL1* in the LS and *tad1* background. Phenotypic analyses demonstrated that RNAi lines with reduced *WL1* expression in the LS background displayed a wide-leaf phenotype, which was significantly increased by 16.1% compared to the LS (Figure 5E-G). Concurrently, the RNAi line with reduced *WL1* expression in the *tad1* background had a wide-leaf phenotype (12.5% wider than that of *tad1*), indicating that the *WL1* RNAi partially rescued the narrow leaf phenotype of *tad1* (Figure 5E-G). These results suggest that TAD1 and WL1 may act, at least in part, in a common pathway to control leaf width.

### WL1 directly represses *NAL1* expression

*WL1* encodes a C2H2 zinc finger protein that contains a conserved C2H2 zinc finger domain and two EAR motifs (Supplemental Figure 5A). Previous studies as well as our results on Y2H revealed that WL1 is a transcription activator (Huang et al., 2009; Li et al., 2013) (Supplemental Figure 5B). Here, a dual-luciferase reporter (DLR) was used to investigate the transcription-regulation activity of WL1. The positive control VP16 showed a relatively high luciferase (LUC) activity whereas the VP16-WL1 fusion protein displayed much lower activity as compared to the positive control (Supplemental Figure 5C). These findings demonstrated that WL1 might also act as a transcriptional repressor.

To identify the target genes of WL1, we used qPCR to analyze the expression of some genes related to regulation of leaf width. Interestingly, the expression of *NAL1* was significantly up-regulated or down-regulated in *wl1* and *tad1*, respectively, as compared to the WT (Figure 6A, B). The contained C2H2 zinc finger domain can bind to a TGNTANN(A/T)T sequence, a cis-element named DST-binding sequence (DBS) (Huang et al., 2009; Li et al., 2013). Sequence analysis revealed the presence of DBS in the promoter region of *NAL1* (Figure 6A). We used chromatin immuno-precipitation (ChIP) assays to test whether WL1 protein could bind to the DBS region of the *NAL1* promoter. Chromatin isolated from young seedlings of the WT was immuno-precipitated with the Anti-WL1 antibody and then subjected to qPCR analysis. The results showed that WL1 protein could bind stably to the F2 site, which contained the DBS (Figure 6C, D). This evidence suggests that WL1 protein might be responsible for the direct regulation of NAL1. Next, we studied the effect of WL1 protein on the expression of the firefly LUC gene reporter, using the DBS-like motif region of *NAL1* as a promoter for transient expression assays in rice protoplasts. Compared with the negative control p35Sm: LUC reporter (Reporter 1), the LUC activity was significantly increased with the pNAL1-35Sm: LUC reporter (Reporter 2). Co-expression of the pNAL1-35Sm: LUC reporter with 35S: WL1 led to a significant decrease of the LUC activity (Figure 6E, F). Therefore, these results indicated that the WL1 protein repressed the expression of *NAL1* by binding the DBS-like motif in the *NAL1* promoter.

**Figure 6.**
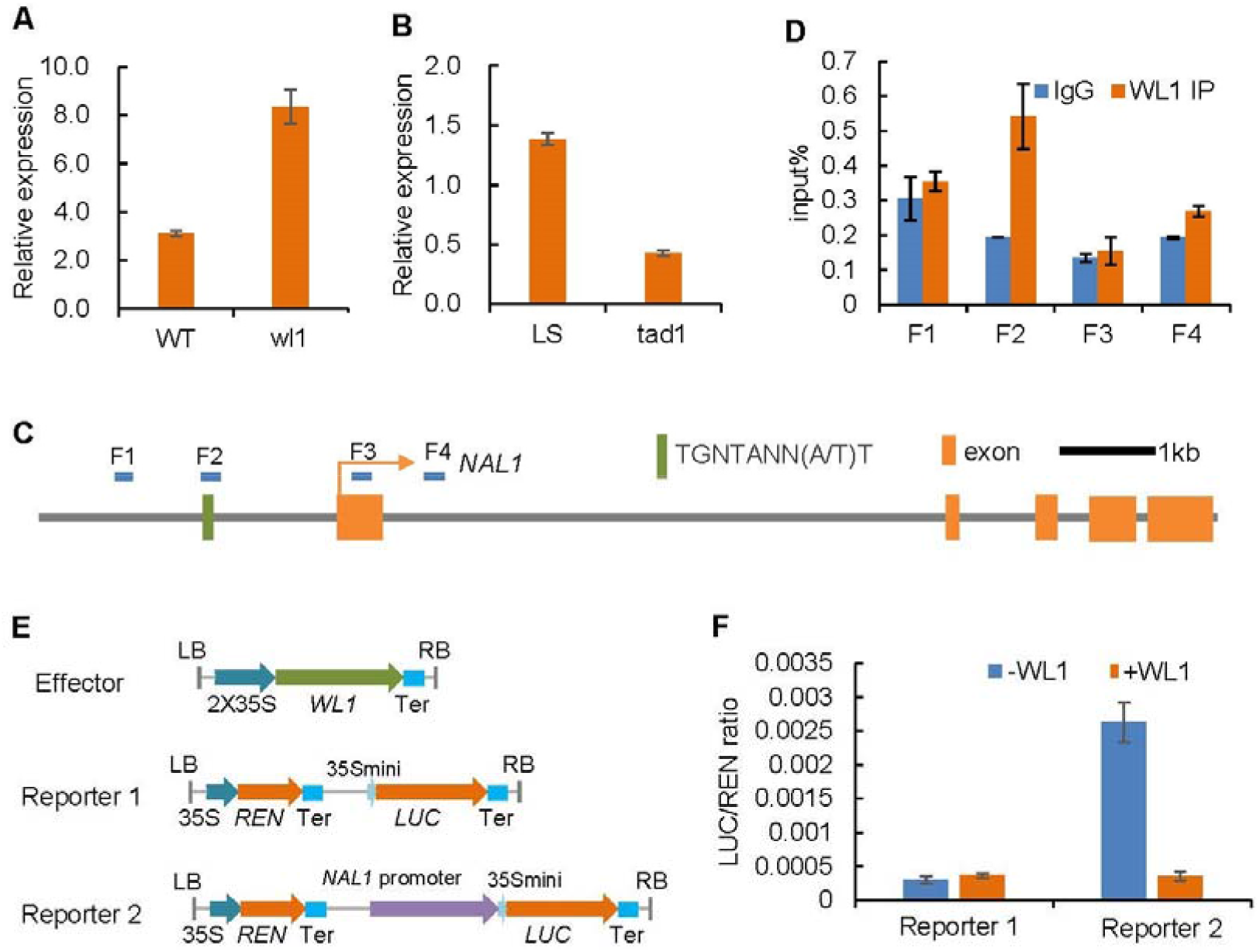
Direct Regulation of *NAL1* Expression by WL1. (A-B) Expression analysis of *NAL1* in leaves of the WT, *wl1* (A) and LS, *tad1* (B) by qRT-PCR. (C) Distribution of potential binding sites in the promoter of *NAL1*. Blue bars represent the DNA fragments amplified in the ChIP assays. (D) ChIP-qPCR for F1-F4 site of *NAL1* with anti-WL1 antibody. ChIP enrichment compared with the input sample was tested by qPCR. (E-F) WL1-repressed *NAL1* expression in vivo. Rice protoplasts transformed with p35Sm:LUC, p35Sm:LUC plus 35S:WL1, pNAL1-35Sm:LUC, pNAL1-35Sm:LUC plus 35S:WL1 or pNAL1-35Sm:LUC plus 35S:WL1. Error bars represent the SD of three repeats.

### WL1 interacts with the OsTPR corepressors to down-regulate the acetylation levels of histone at *NAL1*

The WL1 protein contains two EAR motifs (LxLxL and DLNxxP) at the C- and N-end, respectively (Supplemental Figure 5A). Proteins with EAR motifs are hypothesized to interact with the corepressors TPR. There are three TPR proteins in rice, namely OsTPR1 (LOC_Os01g15020), OsTPR2 (LOC_Os08g06480) and OsTPR3 (LOC_Os03g14980) (Zhuang et al., 2020). To further dissect the molecular mechanism of WL1-mediated regulation of *NAL1* expression, the interaction between WL1 and OsTPRs was examined. First, yeast-two hybrid (Y2H) assays were performed to determine the physical interaction between WL1 and OsTPRs. The results demonstrated that the full-length WL1 protein, the C-terminal EAR motif and the N-terminal EAR motif of WL1 protein as well as the mutated WL1 protein interacted with the three OsTPR proteins in yeast cells (Figure 7A) (Supplemental Figure 6). Second, a bimolecular fluorescence complementation (BiFC) assay was carried out to find out whether the WL1 could interact with OsTPRs in *N. benthamiana* leaves. The yellow fluorescent protein signal was present in the nuclei of epidermal cells of *N. benthamiana* leaves, co-expressing the WL1-cYFP fusion protein with nYFP-OsTPR1, nYFP-OsTPR2, or nYFP-OsTPR3 (Figure 7B). Thus, WL1 interacted with OsTPRs *in vivo*. It is well documented that TPR can interact with histone deacetylases (HDACs) to regulate the acetylation level of histones in the target regions (Zhuang et al., 2020). Therefore, we further detected the histone acetylation level in the *NAL1* by ChIP-qPCR using the chromatin immuno-precipitated by the anti-H3K9ac antibodies. The acetylation levels of H3K9 at P4, P6, P7, and P9-P11 sites, which are largely located in the promoter and first intron of *NAL1*, were significantly increased in the *wl1* mutant compared with those in the WT (Figure 7C, D). These results support WL1 directly binding to the promoter or other regulating regions of the *NAL1* gene. WL1 then recruits the corepressor OsTPRs to inhibit *NAL1* expression by down-regulating the acetylation levels of histones on the chromosomes at *NAL1*.

**Figure 7.**
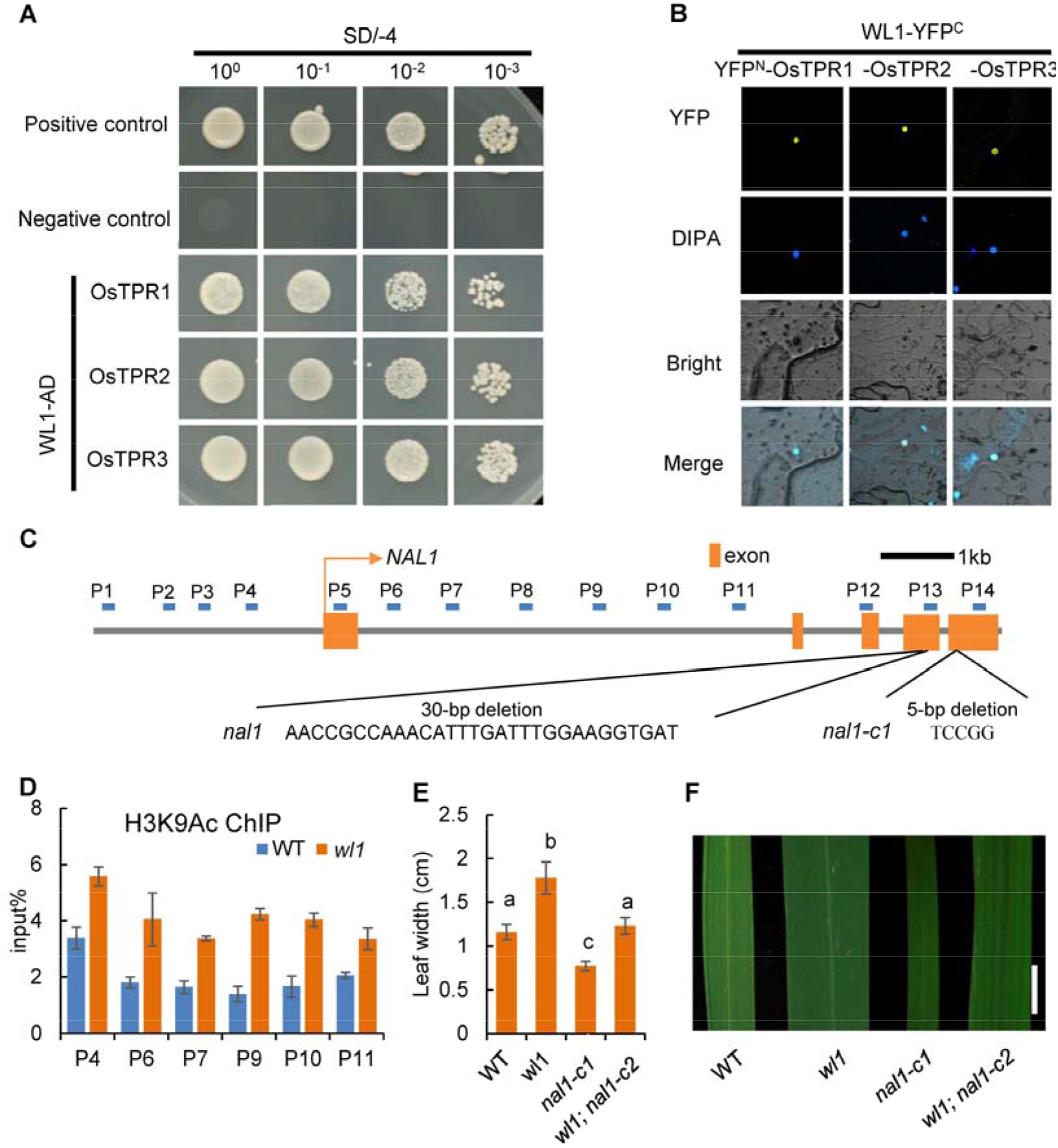
WL1 Interacts with the Corepressors OsTPRs to down-regulate the acetylation levels of histones on the chromosomes at *NAL1*. (A) WL1 interacts with OsTPR1, OsTPR2 and OsTPR3 in yeast cells as shown by Y2H assay. (B) WL1 interacts with OsTPR1, OsTPR2 and OsTPR3 in the nuclei of *N. benthamiana* leaf cells as demonstrated by BiFC assay. (C) Distribution of potential binding sites in the promoter and open reading frame regions of *NAL1* and the mutated sites of *nal1* and *nal1-c1, nal1-c2*. Blue bars represent the DNA fragments amplified in the ChIP assays. (D) ChIP-qPCR for several sites of *NAL1* with anti-H3K9ac antibody in leaves of the WT and *wl1*. (E) Leaf width of the WT, *wl1*, *nal1-c1* and *wl1*; *nal1-c2*. (F) Leaves of the WT, *wl1*, *nal1-c1* and *wl1*; *nal1-c2*. Error bars represent the SD of three repeats. Lowercase letters indicate significant differences (*p*<0.05). Scale bar=1 cm.

To further analyze the genetic interaction between *WL1* and N*AL1*, we generated the loss of function mutant *nal1-c1* using the CRISPR/Cas9 technology in the WT Xinong 1B. The *nal1-c1* had a 5-bp deletion in the CDS of *NAL1*, resulting in a reading frameshift that generated a premature stop codon. A narrow leaf (width reduced by 33.6% compared to the WT) was observed as expected in *nal1-c1* (Figure 7C, E, F). Next, we used the CRISPR/Cas9 technology to knockout *NAL1* in *wl1* background and generate the *wl1*; *nal1-c2* double mutant. The *nal1-c2* had the same 5-bp deletion in *NAL1* as *nal1-c1*. The *wl1*; *nal1-c2* double mutant showed a narrower leaf phenotype than did *wl1* where its leaf width was reduced by 30.9%, indicating that the *nal1-c2* mutation suppressed the leaf width phenotype of *wl1*. In addition, *dst* acted as the allele of *wl1*, also showed width leaf variation, while *nal1* presented narrow leaves. Similarly, the *dst/nal1* double mutant also presented a narrower leaf than in *dst* (Supplemental Figure 7). Overall, the above results suggest that *WL1* and *NAL1* function antagonistically in a common pathway to control leaf width growth in rice (Figure 7E, F).

## Discussion

Leaf morphology is one of the most important components of ideal plant architecture that critically influences photosynthesis in rice. A good leaf shape can optimize the canopy structure, increase the photosynthetic area and ultimately boost grain yield. The molecular mechanism of leaf development has been well-studied in *Arabidopsis thaliana*, which has oval leaves with reticulate veins(Yasui et al., 2018). By contrast, understanding of leaf development in monocots such as cereals, in which the leaves are long and narrow with parallel veins, remains limited despite their agronomic importance. In this study, we isolated a *wide leaf 1* (*wl1*) mutant which exhibited an increased leaf width. We report that the *wl1* phenotype was induced by a mutation in the *DST* gene which encodes a C2H2 zinc finger protein. *DST* regulates drought and salt tolerance as well as grain number per panicle in rice (Huang et al., 2009; Li et al., 2013; Cui et al., 2015). However, how *DST* participates in the regulation of leaf width has not been thoroughly investigated. It is also a great future challenge to investigate whether *DST* could coordinate stress responses, grain size, and leaf width.

To elucidate the involvement of *WL1* in the control of leaf width, we demonstrated that the APC/C^TAD1^-WL1-NAL1 pathway promoted leaf growth, suggesting that this pathway holds a potential to increase leaf width in key crop plants. TAD1 is an activator of the anaphase-promoting complex/cyclosome (APC/C) complex and regulates the development of tillers and roots (Lin et al., 2012; Xu et al., 2012; Lin et al., 2020). In this study, loss of function of *TAD1* caused the narrow leaf phenotype, indicating that *TAD1* is a positive regulator of leaf width in rice. The *wl1* mutant and RNAi lines showed altered leaf width, indicating that *WL1* is a negative regulator of this phenotype. Biochemical analyses indicated that TAD1 interacts with and mediates the degradation of WL1 via the ubiquitin-26S proteasome pathway. Additionally, genetic analyses suggested that TAD1 and WL1 act in a common pathway to regulate leaf width. Overall, we hypothesize that TAD1 and WL1 act as a module to regulate leaf width. Furthermore, we provide evidence that WL1 directly binds to the promoter or other regulating regions of the *NAL1* gene and then recruits the corepressor OsTPRs. These repressors inhibit *NAL1* expression by down-regulating the acetylation levels of histone on chromosomes at *NAL1*, indicating that *NAL1* is a target gene for WL1. Taken together, our genetic and biochemical analyses revealed that APC/C^TAD1^-WL1-NAL1 function in a positive regulatory pathway for leaf width in rice. Based on these results, we propose a working model of how the APC/C^TAD1^-WL1-NAL1 module controls leaf width in rice (Figure 8). Simply, TAD1 activates the APC/C E3 ubiquitin ligase which targets WL1 for degradation; WL1 negatively regulates the expression of *NAL1*, and consequently represses leaf width (Figure 8).

**Figure 8.**
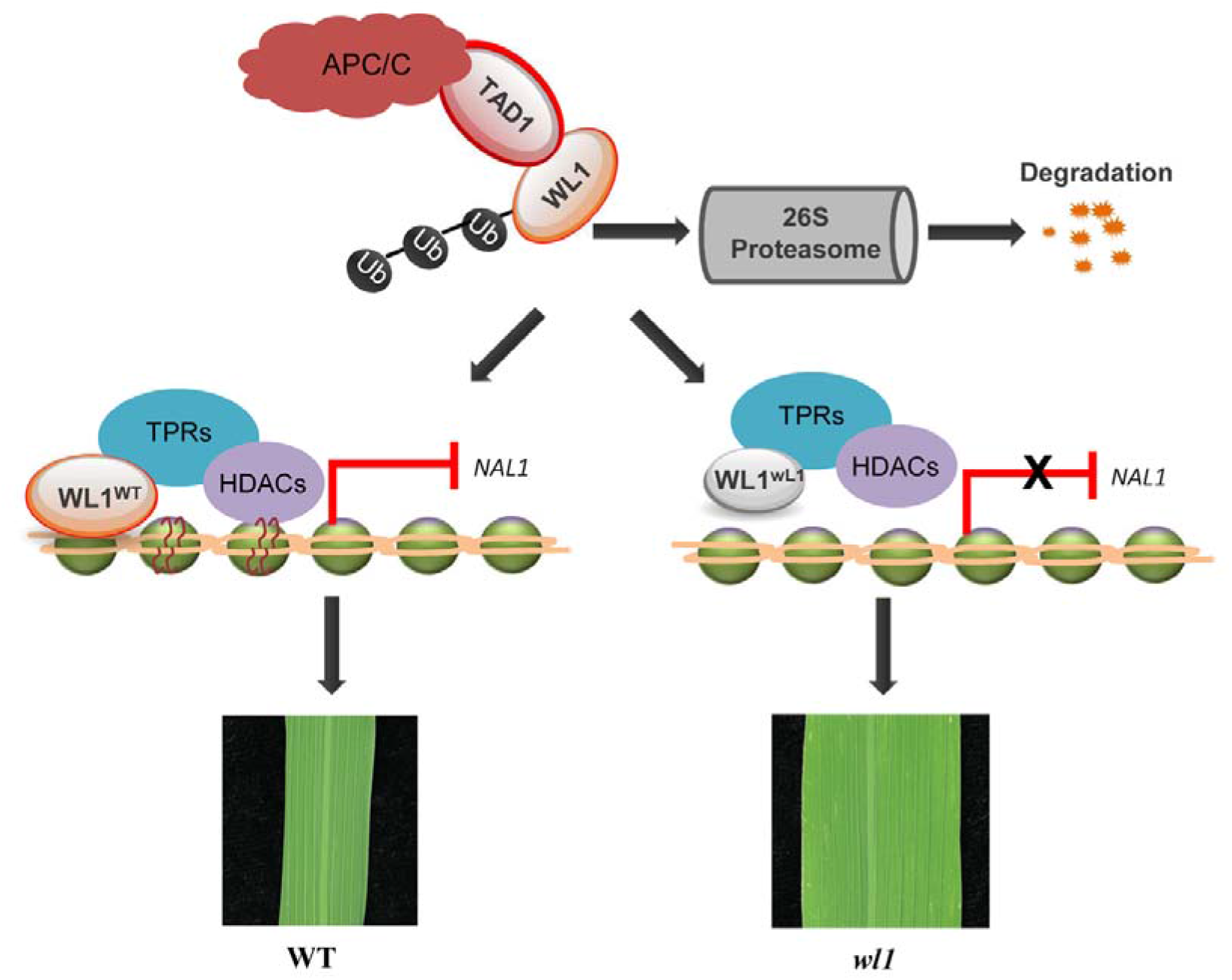
A proposed model for APC/C^TAD1^-WL1-NAL1 module-mediated control of leaf width. TAD1 activates the APC/C E3 ubiquitin ligase activity and targets WL1 for degradation. WL1 negatively regulates the expression of *NAL1*, and consequently decreases leaf width.

Previous studies have identified a variety of narrow leaf mutants in rice including the auxin-related mutants *nal1, fib, nal7*, and *tdd1*(Fujino et al., 2008b; Qi et al., 2008b; Sazuka et al., 2009; Yoshikawa et al., 2014); the cellulose synthase-like D (CSLD)-related mutant *nrl1*(Li et al., 2009) and the polarity-related mutants *sil1* and *lf1*(Zhang et al., 2009; Zhang et al., 2021). Therefore, the molecular mechanisms of the narrow leaf trait have been extensively studied. However, studies on mutants with wide leaf traits along with the involved molecular mechanisms are scarce. The *D62* gene encodes a protein of the GRAS family, which are inhibitors of GA signaling. Mutations in this gene can produce dwarf plants with more erect, wider, shorter, and darker leaves (Tong et al., 2009). *Osa-MIR319a* and *Osa-MIR319b* belong to the *miR319* gene family. Overexpression of *Osa-MIR319a* and *Osa-MIR319b* in rice leads to increasing the leaf width. The numbers of small vascular bundles in the *osa-miR319*-overexpressing plants was significantly increased compared with those in the WT, indicating that the wide leaves of the *osa-miR319*-trangenic plants resulted from increasing the numbers of small vascular bundles and numbers of cells between them (Yang et al., 2013). Here, we identified a novel wide-leaf mutant with significant increase in the area and width of leaf primordia, the distances between the small vascular bundles and the numbers of small and large vascular bundles as compared to the WT. Therefore, *WL1* is likely to play an important role in regulation of leaf width development by controlling the numbers of large vascular bundles and the distances between small vascular bundles. In addition to the altered leaf width, the *wl1* mutant displayed a larger panicle with a significantly increased numbers of grains per panicle (Supplemental Figure 7). Thus, we propose *WL1* as a prime candidate for further studies to support the genetic improvement of rice leaf phenotypes and grain production.

## Materials and methods

### Plant materials

The rice mutant *wl1* was isolated from an ethyl methane sulfonate-mutagenizd population of *O. sativa* subsp. *indica* cultivar Xinong 1B. Xinong 1B was used as the wild-type (WT) reference for phenotypic observation and *in situ* hybridization analyses. All plant materials were grown in the experimental fields of the Rice Research Institute of Southwest University in Chongqing and Hainan, China.

### Map-based cloning of *WL1*

The *wl1* mutant was crossed with ‘Jinhui 10’ (bred by the Rice Research Institute, Southwest University), and 544 F_2_ plants exhibiting the mutant phenotype were selected for gene mapping. Gene mapping was conducted using simple sequence repeat markers obtained from the publicly available rice databases Gramene (http://www.gramene.org) and the Rice Genomic Research Program (http://rgp.dna.affrc.go.jp/E/publicdata/caps/index.html). The insertion/deletion markers were developed by comparing the genomic sequences of ‘Xinong1B’ to those of ‘Jinhui 10’ in our laboratory. The primer sequences used in gene mapping and candidate gene analyses are provided in Supplementary Table S1.

### Vector construction for plant transformation

For the complementation test, a 3776-bp genomic fragment was cloned into the binary vector pCAMBIA1301. The resulting recombinant plasmids were introduced into *wl1* using the *Agrobacterium tumefaciens*-mediated transformation method as described previously (Zhang et al., 2017a). To generate a construct for RNAi, a 264-bp fragment of the WL1 complementary DNA was amplified and inserted into the vector pTCK303 to obtain the intermediate vector. The resulting recombinant plasmids were transformed into the WT (Xinong 1B), LS, and *tad1* mutant, respectively, using the *Agrobacterium-mediated* transformation method (Zhang et al., 2017a). For the knockout assay, the 20-bp target sequence (5’TGATGCTCCTCGGAGCGTCT3’) from the *NAL1* genomic fragment for sgRNA targeting was cloned into the CRISPR/cas9 expression vector to generate the CRISPR/cas9-mediated construct pCRISPR-*NAL1* (Ma et al., 2015). The resulting recombinant plasmid was introduced into the WT (Xinong 1B), *wl1* mutants, respectively, using the *A. tumefaciens*-mediated transformation method (Zhang et al., 2017a). The used primer sequences are listed in Supplementary Table S1.

### RNA isolation and qRT-PCR analysis

Total RNA of rice was isolated from the roots, stems, leaf sheaths, leaves, and young panicles using the RNAprep Pure Plant RNA Purification Kit (Tiangen, Beijing, China). First-strand complementary DNA was synthesized from 2 μg total RNA using oligo(dT)_18_ primers in a 25 μL reaction volume with the SuperScript III Reverse Transcriptase Kit (Invitrogen, Carlsbad, CA, USA). The reverse-transcribed RNA (cDNA) (0.5 μL) was used as a template for PCR amplification with gene-specific primers (Supplemental Table S1). Quantitative PCR (qPCR) analysis was performed with the CFX Connect™ Real-Time System (Bio-Rad, Berkeley, CA, USA) and SYBR permix Ex Taq II Kit (TaKaRa, Kyoto, Japan); using *actin* as an internal control. A minimum of three replicates was analyzed to produce the mean values of the expression level of each gene.

### Microscopic analysis

The aerial tissues containing SAMs and leaf primordia from *wl1* and WT were collected and immediately fixed in FAA solution (50% ethanol, 0.9 M glacial acetic acid, and 3.7% formaldehyde) and maintained at 4°C overnight. The samples were dehydrated in a series of ethanol solutions, substituted with xylene and embedded in paraffin (Sigma, St. Louis, MO, USA). The samples were sectioned to 8 μm thickness, mounted onto poly-L-Lys-coated glass slides, de-paraffinized with xylene, and dehydrated through a series of ethanol solutions. The sections were stained with 1% safranine (Amresco, Framingham, MA, USA) and 1% Fast Green (Amresco), dehydrated through a series of ethanol solutions, cleared with xylene and made permanent by adding a coverslip. Light microscopy was performed using the Eclipse E600 microscope (Nikon, Tokyo, Japan).

### Multiple sequence alignment and construction of phylogenetic tree

Protein sequences were obtained with BLAST (Basic Local Alignment Search Tool) in PHYTOZOME, using an EXPECT value threshold of 10^-5^ (http://phytozome.jgi.doe.gov/pz/portal.html#!search?show=BLAST). A phylogenetic tree was constructed using MEGA 5.0 (Tamura et al., 2011). The tree was constructed using the maximum likelihood method based on the Jones, Taylor, Thornton matrix-based model with the lowest Bayesian information criteria scores (Tamura et al., 2011). Each node’s bootstrap support values from 500 replicates were included next to the branches.

### Subcellular localization

The full-length coding region of *WL1* was amplified and cloned into the expression cassette 35S-*GFP*-NOS (pAN580) to generate *GFP-WL1* and *WL1-GFP* fusion vectors. Then, *GFP, GFP-WL1*, and *WL1-GFP* plasmids were inserted into rice protoplasts as previously described (Zhang et al., 2017b). After overnight incubation at 28°C, GFP fluorescence was observed with a confocal laser scanning microscope (Olympus, Tokyo, Japan). The used primers are listed in Supplemental Table S1.

### *In situ* hybridization

For WL1 probe, gene-specific cDNA was amplified and labeled using the DIG RNA Labeling Kit (Roche, Basel, Switzerland). Pretreatment of sections, hybridization, and immunological detection was performed as described previously (Zhang et al., 2017a). The primers used are listed in Supplemental Table S1.

### Transcriptional activation assay

The transcriptional activity of WL1 in rice protoplasts was analyzed using the dual-luciferase reporter (DLR) assay system. The DLR assay system was used with the GloMax 20-20 luminometer (Promega, Madison, WI, USA) to measure the relative luciferase (LUC) activity (Zhuang et al., 2020). The full length and truncated coding frames of *WL1* were fused to the *GAL4* DNA-binding domain (BD) driven by the 35S promoter. The transcriptional activator VP16 was used as a positive control whereas GAL4-BD was the negative control. VP16, BD-WL1, VP16-WL1, and GAL4-BD effectors were transiently expressed in rice protoplasts. The primers used are listed in Supplemental Table S1.

### Chromatin immuno-precipitation-qPCR

The WT and *wl1* plants were subjected to the ChIP analysis. Panicles were collected for isolation of nuclear extracts. The EpiQuik Plant ChIP Kit (P-2014-48, Epigentek), anti-WL1 antibody, and anti-Histone H3 (acetyl K9) antibody (ChIP grade; ab10812, Abcam) were used in the ChIP assays. Immunoglobulin G (IgG) antibody was included as a negative control. All PCR experiments were conducted using 40 cycles of 95°C for 5 s, 60°C for 30 s, and 72°C for 30 s. The reaction mixtures contained 10 pmol of each primer and 1 mL of DNA from either ChIP, control or input DNA diluted 20-fold (per biological replicate) as a template. A minimum of three biological repeats (1 g of panicle sample), each with three technical repeats, was analyzed to produce data for statistical analysis. Experimental procedures for ChIP-qPCR were performed as previously described previously (Zhang et al., 2017a). The primer sequences are listed in Supplemental Table 1.

### Transient expression assay

The *NAL1* promoter and a mini cauliflower mosaic virus 35S promoter were amplified and cloned into the pGreenII0800-LUC double-reporter vector. To generate the 2×35Spro:: *WL1* constructs, the full-length coding region of WL1 was amplified with PCR and recombined into the pAN580 vector, including two CaMV35S promoters. The constructs were then introduced into rice protoplasts as previously described (Zhang et al., 2021). After overnight incubation at 28°C, LUC and Renilla (REN) luciferase activities were measured using the Dual Luciferase Assay Kit (Promega) and analyzed using the Luminoskan Ascent Microplate Luminometer (Thermo Fisher Scientific, Waltham, MA USA) according to the manufacturer’s instructions. The results were calculated from the ratios of LUC/REN (Zhuang et al., 2020). A minimum of six transient assay measurements was carried out for each sample. The primers used are listed in Supplemental Table S1.

### Yeast two-hybrid assay (Y2H)

The Y2H assays were performed using the Matchmaker Gold Yeast Two-Hybrid System (Clontech, Mountain View, CA, USA). The full-length and truncated coding regions of *WL1* were amplified and cloned into the yeast expression vector pGADT7 (Clontech) to produce pGADT7-WL1, pGADT7-N-WL1, and pGADT7-C-WL1. The full-length coding regions of *TAD1*, *OsTPR1, OsTPR2,* and *OsTPR3* were introduced into the vector pGBKT7 (Clontech) to produce pGBKT7-TAD1, pGBKT7-OsTPR1, pGBKT7-OsTPR2, and pGBKT7-OsTPR3. The Y2HGOLD yeast strain was used in the Y2H assays. The pGADT7-T1pGBKT7-lam served as a negative control and pGADT7-T1pGBKT7-53 as a positive control. According to the manufacturer’s instructions, these plasmids were co-transformed into the Y2H GOLD strain in an activation domain (AD)-BD coupled manner (Yeast Protocols Handbook, PT3024-1; Clontech). The primers used are listed in Supplemental Table S1.

### Bimolecular fluorescence assay

The vectors pXY104 and pXY106 harboring the fragments encoding the C- and N-terminal halves of YFP (cYFP and nYFP), respectively were used to produce constructs for the bimolecular fluorescence assays (BiFC). The cDNA fragment encoding WL1 was fused to the fragment encoding the C-terminus of YFP, whereas the cDNA fragments encoding *TAD1, OsTPR1, OsTPR2* or *OsTPR3* were fused to the fragment encoding the N-terminus of YFP. All vectors were introduced into the *Agrobacterium* strain GV3101. The *Agrobacterium* cultures harboring the constructs expressing nYFP and cYFP fusion proteins were mixed in a ratio of 1:1 and introduced into tobacco leaves by infiltration. The BiFC signals were detected by confocal microscopy as described above. The primers used are listed in Supplemental Table S1.

### Pull-down assay using purified proteins

The pull-down assays were performed as previously described with minor modifications (Zhang et al., 2021). The *WL1* CDS fragments were cloned into the pGEX-4T-1 vector whereas *TAD1* cDNA fragments were cloned into pET32a. GST-WL1 and His-TAD1 proteins were expressed in *E. coli* (BL21). His-TAD1 fusion proteins were purified according to the manufacturer’s protocol (New England Biolabs, Ipswich, MA, USA). The bait protein GST-WL1 was incubated with GST beads (GE Healthcare, Chicago, IL, USA) in non-EDTA buffer (20 mM Tris-HCl, pH 7.5 and 100 mM NaCl) at 4°C for 2 h and then washed three times with the same buffer. The beads were re-suspended in 1 mL non-EDTA buffer before 50-100 ng His-TAD1 prey protein was added to the solution. The mixture was then incubated at 4°C for 1 h and rinsed three times. Proteins were eluted with sodium dodecyl sulfate loading buffer and subjected to immuno-blot analysis. The prey and bait proteins were detected using anti-His (1:1000, No. #2365; Cell Signaling Technology [CST], Danvers, MA, USA) and anti-GST (1:1000, No. #2622; CST) antibodies, respectively. The primers used are listed in Supplemental Table S1.

### Antibody preparation and western blot analysis

The CDS of *WL1* was inserted into the pGEX-4T-1 vector. The purified GST-WL1 fusion protein was injected into rabbits to produce polyclonal antibodies for WL1. Total protein extracts from the WT, *wl1*, LS, and *tad1* young panicles were isolated using a Plant Total Protein Extraction Kit (Sangon Biotech). Western blots were performed using primary antibodies against WL1 and ACTIN (K101527P, Solarbio, 1:2000 dilution). The AP-conjugated Affinipure Goat Anti-Rabbit IgG (Jackson ImmunoResearch) was used as secondary antibody according to the recommended dilution ratios in the iBind Cards, using the iBind Western Device (Invitrogen) and detected using the NBT/BCIP substrate (Sangon Biotech).

### Cell-free degradation assay

The cell-free degradation assay was performed as reported previously (Lin et al., 2012; Xu et al., 2012). The total proteins from *tad1* and WT seedlings were extracted with the degradation buffer (25 mM Tris-HCl, pH 7.5, 10 mM NaCl, 10 mM MgCl_2_, 4 mM PMSF, 5 mM dithiothreitol, and 10 mM ATP) and adjusted to equal concentrations with the degradation buffer. The MG132 (T510313-0001, Sangon Biotech) was selectively added to various *in vitro* degradation assays as indicated. To degrade purified GST and GST-WL1 proteins, equal amounts of the purified proteins (about 100 ng) were incubated in 100 μl aliquots of rice total-protein extracts (containing about 500 μg total proteins) at 28°C for each assay. An equal amount of solvent for each reagent was used for the mock control. The extracts were then incubated at 28°C and the samples were collected at the indicated intervals for determination of GST and GST-WL1 protein abundance by western blots using the anti-GST (#2625, Cell Signaling Technology, 1:1000 dilution) and alkaline phosphatase-conjugated anti-rabbit (# 31340, Thermo Fisher Scientific, 1:5000 dilution). The chemi-luminescence detection was performed with DIG Substrate (Roche, Basel, Switzerland).

### *In vitro* ubiquitination assay

The *in vitro* ubiquitination assay was performed as reported previously (Lin et al., 2012; Xu et al., 2012). The purified GST-WL1 protein bound to the Glutathione Sepharose (P2020, Solarbio) was incubated at 28°C with equal amounts of the crude extracts of rice seedlings in a buffer containing 25 mM Tris-HCl, pH 7.5, 10 mM MgCl_2_, 5 mM dithiothreitol, 10 mM NaCl, 10 mM adenosine triphosphate, and 40 μM MG132. After incubation at 28°C for the indicated intervals, the GST-WL1 and GST-WL1-(Ub)n fusion proteins, dissolved in SDS-PAGE loading buffer, were added. The samples were then loaded onto SDS-PAGE gels. Western blots were performed with antibodies against the WL1.

## Acknowledgments

We thank Jiayang Li (Chinese Academy of Sciences) for kindly providing the LS and *tad1* seeds. We also thank Chuanyou Li (Chinese Academy of Sciences) for kindly providing the ZF802, *nal1, dst*, and *dst/nal1* seeds. Additionally, we are grateful to Weixun Wu (China National Rice Research Institute) for providing the CRISPR/cas9 expression vector. This work was supported by the National Natural Science Foundation of China, Grants 31900612 and 31730063.

## Author Contributions

T.Z. and G.-H.H. conceived and designed the experiments; J.Y., W.-W.X., Y.Z., L.Y., P.Y., G.-L.Y., X.-F.Z., Z.-F.H., Y.X. conducted the experiments; T.Z., J.Y., W.-W.X., X.-C.S., Y.-F.L., F.-M.Z., Y.-H.L. analyzed the data; and T.Z., G.-H.H., J.Y wrote the manuscript.

